# Augmenting CAR NK cell Anti-tumor Activity by Synapse Tuning

**DOI:** 10.1101/2022.10.26.512898

**Authors:** Peter J Chockley, Jorge Ibanez-Vega, Giedre Krenciute, Lindsay J Talbot, Stephen Gottschalk

## Abstract

Chimeric antigen receptor (CAR) technologies have been clinically implemented for the treatment of hematological malignancies; however, solid tumors remain resilient to CAR therapeutics^1-3^. Natural Killer (NK) cells may provide an optimal class of immune cells for CAR-based approaches due to their inherent anti-tumor functionality. We sought to tune CAR synapses in NK cells by adding an intracellular scaffolding protein binding site to the CAR. We employed a PDZ binding motif (PDZbm) that specifically binds Scribble^4^ resulting in additional scaffolding crosslinking to enhance synapse formation and cell polarization^5,6^. Combined effects of this novel CAR design resulted in increased effector cell functionality *in vitro* and *in vivo*. Synapse-tuned CAR-NK cells exhibited amplified synaptic strength, number and abundance of secreted cytokines, enhanced killing of tumor cells, and prolonged survival with tumor clearance in two solid tumor models. Thus, synapse tuning has the potential to improve the efficacy of CAR-based cell therapeutics.

## Main

CAR technology has been utilized to elicit antigen-dependent signaling pathways in both NK and T cells^7^. However, CAR:Antigen complexes form disordered synapses^8^, which do not consist of bona fide central, peripheral, or distal supramolecular activation complexes (SMACs) in contrast to canonical T cell Receptor:Human Leukocyte Antigen (TCR:HLA) synapses^8^. These organized SMACs allow for a lower threshold for antigen recognition^9^. Further, the additional signaling molecules recruited to the synapse increase the efficiency of signaling and exclude inhibitory phosphatases^10^. In contrast, the disjointed CAR synapse is a punctate structure with islands of CAR:Antigen complexes^9,11^. These islands are, putatively, open to dephosphorylation and thus require a larger number of interactions to initiate downstream signaling and activate effector cells. We theorized that synapse modulation or tuning to an increasingly ordered state would result in an efficient and effective CAR. We focused our work on NK cells since they have an innate ability to kill tumor cells and contain ∼5-to 7-fold more lytic granules compared to T cells^12,13^. Likewise, NK cells are not known to cause graft versus host disease and tumor cells have a reduced ability to evade attack due to the multiple targeting paradigms that are employed outside of the CAR:Antigen recognition axis^14^.

To improve CAR synapse formation, we focused on Postsynaptic density-95, Discs large, and Zona occuldens 1 binding moieties (PDZbms). There are roughly 400 proteins that contain PDZ domains, many of these are typically associated with highly polarized epithelial, endothelial, and neuronal cells^15^. Additionally, there are some PDZbms that aid in the synapse formation and polarity of immune cells^4^. One of these, Cytotoxic and Regulatory T cell Associated Molecule (CRTAM) is a binding partner of Scribble, a PDZ domain containing intracellular scaffolding protein and mainly described in epithelial junctions^5^. CRTAM to Scribble interactions have been studied in T and NK cells^4,16^, and CRTAM plays a role in NK cell tumor immunosurveillance^17^. Thus, we selected the PDZbm of CRTAM and added it to the C-terminus (CAR.PDZ) of a CD28z CAR (CAR)^18^, which targets the tyrosine kinase, ephrin type-A receptor 2 (EphA2), a tumor associated antigen expressed in a broad range of solid tumors^19,20^ (**Figure 1a**). CAR NK cells were generated from cultured primary peripheral blood NK cells by retroviral transduction and CAR, CAR.PDZ, and a non-functional control CAR (CAR Δ) were expressed equally with no differences on the cell surface of NK cells across numerous donors (**Extended Data Figure 1a,b**). Additionally, the CAR modifications did not alter the immunophenotype of NK cells (**Extended Data Figure 1c**).

**Figure 1:**
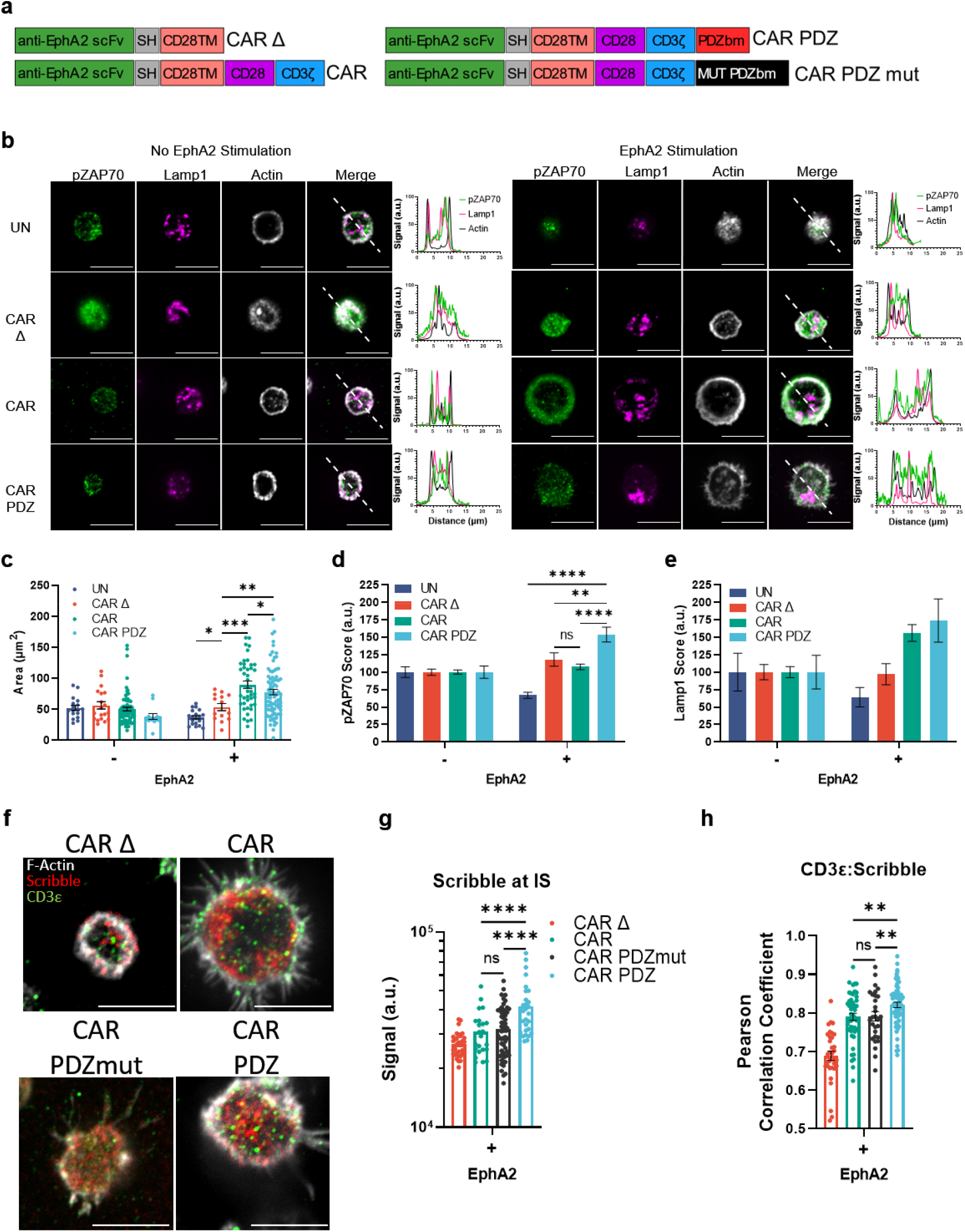
The PDZ binding moiety scaffolding anchor enhances CAR NK cell synapse formation. (a) Chimeric antigen receptor design schemes. Antigen recognition domain (anti-EphA2 scFv): green, short hinge domain IgG (SH): grey, transmembrane domain CD28 (CD28TM): salmon, CD28 co-stimulatory domain: purple, CD3ζ activation domain: blue, PDZbm scaffolding anchor domain: red, and mutated PDZbm domain in black (T2A.truncated CD19 tag of the retroviral vector not shown). (b) Fluorescent confocal microscopy of NK cells plated on a poly-L-lysine coated slide with and without recombinant human EphA2 protein incubated for 30 minutes, white bars indicate 10 microns; pZAP70 (green), Lamp1 (magenta), Actin (white). Dashed line in merged images delineate the approximate line scan quantization of the fluorescent signal depicted in the histograms to the right. Quantified single cell fluorescent data is plotted in (c-e). (c) Quantification of synaptic area determined by actin immunolabeling, n=17 and 19, UN; n=19 and 16, CAR Δ; n=76 and 44, CAR; n=12 and 83, CAR.PDZ at 0 and 30 minutes respectively. Two-Way ANOVA was used to determine statistical significance with Two-stage linear step-up procedure of Benjamini, Krieger and Yekutieli to correct for FDR. Statistical difference delineated by q<0.05 *, q<0.01 **, q<0.001 ***; mean±SEM shown. (d) pZAP70 intensity quantification from (c). Arbitrary Units (a.u.), defined by the intensity of the fluorescent signal per unit area of the synapse (actin ring). All cells were normalized to the mean of 0 minutes. Two-Way ANOVA was used to determine statistical significance with Two-stage linear step-up procedure of Benjamini, Krieger and Yekutieli to correct for FDR. Statistical difference delineated by q<0.01 **, q<0.0001 ****; mean±SEM shown. Cells are the same as in (c). (e) Lamp1 intensity quantification. a.u. defined by the intensity of the fluorescent signal per unit area of the synapse (actin ring). All cells were normalized to the mean of no EphA2; mean±SEM shown. Cells are the same as in (c). (f) Confocal images as prepared in (b) incubated for 60minutes with NK cells in various groups quantified in (g,h). White bars indicate 10microns. Immunolabelling of Scribble in red, CD3ε in green, and filamentous actin (F-actin) in white. (g) Scribble polarization and accumulation at the immune synapse (IS). CAR Δ; n=32, CAR; n=23, CAR.PDZmut: n=57, CAR.PDZ n=28 from two independent experiments, One-Way ANOVA was used to determine statistical significance with Two-stage linear step-up procedure of Benjamini, Krieger, and Yekutieli to correct for FDR. Statistical difference delineated by q<0.01 **, q<0.001 ***; mean±SEM shown. (h) CD3ε co-localization with Scribble as determined by Pearson Correlation Coefficient. CAR Δ; n=33, CAR; n=48, CAR.PDZmut: n=29, CAR.PDZ n=61, One-Way ANOVA was used to determine statistical significance with Two-stage linear step-up procedure of Benjamini, Krieger, and Yekutieli to correct for FDR. Statistical difference delineated by q<0.05 *, q<0.0001 ****; mean±SEM shown.

We first determined if the PDZbm modulates CAR NK cell synapse formation in the singular context of EphA2 by doping recombinant human EphA2 protein to poly-L-lysine coated glass slides and allowed CAR NK cells to incubate and interact for 30 minutes (**Figure 1b**). We assessed synaptic area, downstream signaling via ZAP70 phosphorylation, and lysosomal polarization via confocal microscopy. Representative images, selected by the corresponding mean value calculated, are taken at the z-plane just above the glass surface to acquire a circular cross section of the spheroidal NK cells as they rest on top interacting with the EphA2 protein. We found a marked difference between CAR.PDZ and CAR constructs in the synaptic area (**Figure 1c**) and pZAP70 accumulation at the immune synapse (**Figure 1d**). The synaptic area was more condensed for CAR.PDZs and we observed that pZAP70 was increased in CAR.PDZ NK cells suggesting a more efficient signaling cascade (**Figure 1d**). Lysosomal polarization was not statistically different; however, it trended toward elevated levels in CAR.PDZ NK cells, which suggests cytotoxic vesicle recruitment to the immune synapse is enhanced (**Figure 1e**). We also found that CAR.PDZ constructs induced a significantly greater accumulation of Scribble (**Figure 1f,g**), confirming the functionality of the PDZ domain. This accumulation was time dependent as differences were not observed at 30 minutes (**Extended Figure 2a,b**). Furthermore, we observed an increase in a cytoplasmic splice variant of CD3ε^21^ that co-localized with Scribble as determined by the Pearson correlation coefficient specifically in CAR.PDZ NK cells (**Figure 1h**).

We next sought to investigate other synaptic protein polarizations. Given the importance of f-actin polymerization in synapse formation we elected to explore Wiskott-Aldrich syndrome protein (WASp). WASp regulates f-actin polymerization^22^ and we found a rapid accumulation of WASp in PDZ.CAR NK cells upon antigen recognition (**Extended Figure 3a**). Thus, including a PDZ domain results in improved synapse formation, as judged by a condensed synaptic area with higher levels of phosphorylation (pZAP70) and recruitment of additional signaling molecules (CD3ε).

Given the multifactorial signal integration that NK cells calculate on a per cell basis^14,23^, we opted to explore the complex binding between NK cells and tumor cells, specifically EphA2 expressing A549 lung adenocarcinoma cells. We utilized a single cell avidity measurement technology, z-MOVI™, that determines the precise binding force between effector and target cell. We show that CAR.PDZ constructs have amplified binding capabilities after a 5 minute co-culture period followed by acoustic force exposure (**Figure 2a,b**). Intriguingly, we found no difference between standard CAR and non-signaling CAR constructs which further demonstrates the lack of internal super structure being formed in a traditional CAR signaling cascade (**Figure 2c**). CAR binding specificity was confirmed with an A549 cell line in which EphA2 was knocked out (KO) by CRISPR/Cas 9 gene editing (**Figure 2d,e**). We expanded upon these findings with another cancer cell line, LM7 and found similar results (**Extended Data Figure 4a**). Additionally, we created another CAR containing the PDZbm domain targeting the solid tumor antigen B7-H3 (**Extended Data Figure 5a,b**),^24,25^ and found similar results (**Extended Data Figure 5c**).

**Figure 2:**
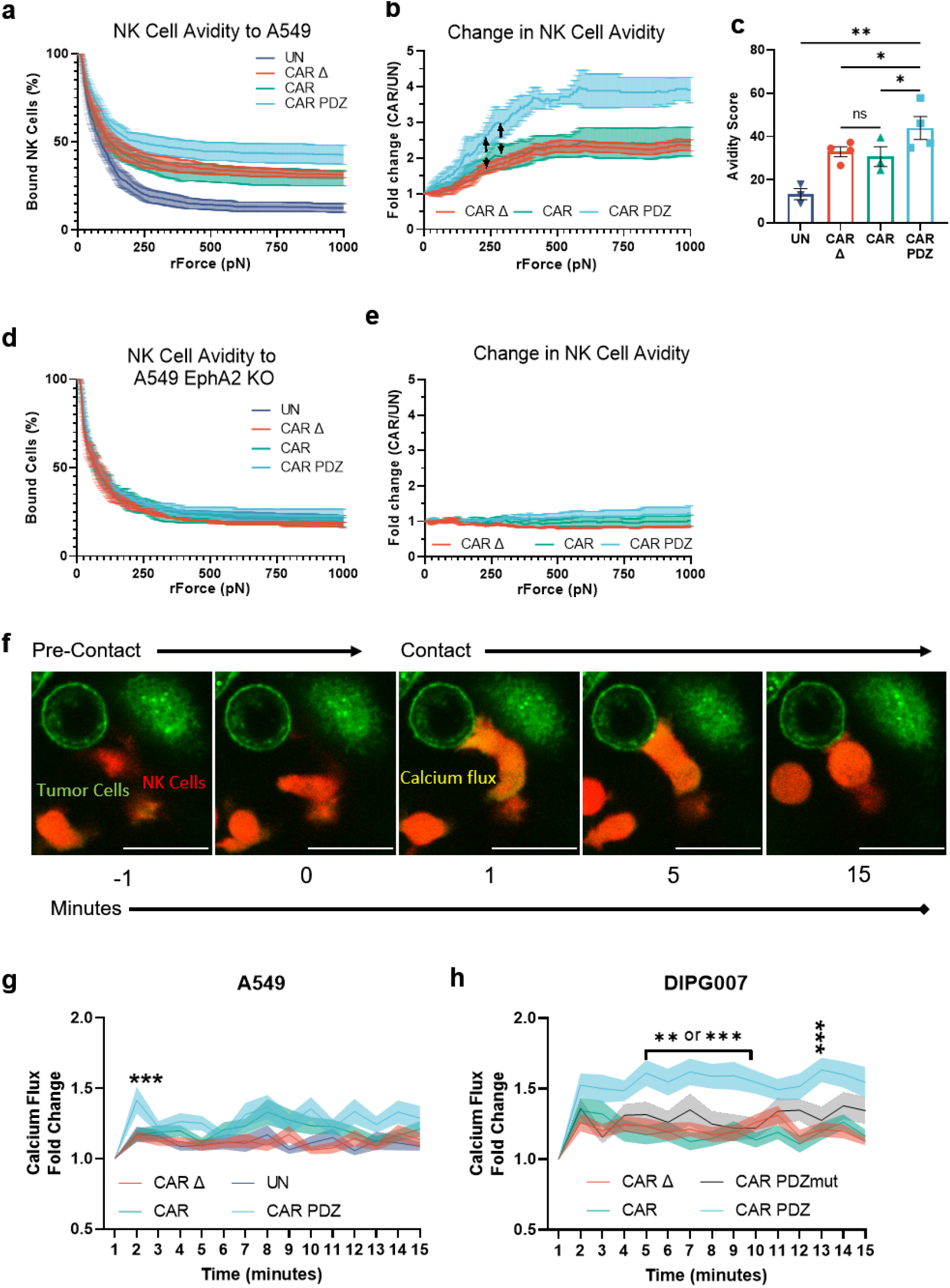
CAR.PDZ NK cells exhibit enhanced avidity and calcium flux upon cancer cell recognition. (a) Single cell assessment of NK cell avidity to EphA2-positive A549 tumor cells after 5 minutes of co-culture; n>100 for each independent experiment, 2-4 merged independent experiments per condition; mean±SEM shown. (b) Normalized fold change of CAR NK cell binding compared to untransduced NK cells from (f). Arrowed lines indicate the point of statistical difference at CAR.PDZ vs. CAR Δ at 231pN and CAR.PDZ vs CAR at 251pN which continued to 1000pN. Two-Way ANOVA was used to determine statistical significance with Two-stage linear step-up procedure of Benjamini, Krieger and Yekutieli to correct for FDR; mean±SEM shown. (c) Avidity score of CAR NK cells determined by plateau of One-Phase Decay analysis from (f). One-Way ANOVA was used to determine statistical significance with Two-stage linear step-up procedure of Benjamini, Krieger and Yekutieli to correct for FDR. Statistical difference delineated by q<0.05 *, q<0.01 **; mean±SEM shown. (d) Same experimental conditions as (a) except utilizing EphA2 deleted A549 cells. (e) Same normalized fold change as described in (c) utilizing EphA2 deleted A549 cells. (f) Representative single cell calcium flux analysis of NK cells interacting with A549 tumor cells over indicated time points. Tumor cells labelled in green, NK cells in red, and calcium flux seen as a yellow burst. (g) Calcium flux quantification of NK cell interactions with A549 tumor cells over 15 minutes. UN; n= 28, CAR Δ; n=35, CAR; n=44, CAR.PDZ n=48. Two-Way ANOVA was used to determine statistical significance with Two-stage linear step-up procedure of Benjamini, Krieger and Yekutieli to correct for FDR; mean±SEM shown. q<0.001 ***; mean±SEM shown. (h) Calcium flux quantification of NK cell interactions with DIPG007 tumor cells over 15 minutes. CAR.PDZ significance from all other groups were seen from 5 to 10 minutes and at 13 minutes. CAR Δ; n=30, CAR; n=19, CAR.PDZmut: n=26, CAR.PDZ n=43, Two-Way ANOVA was used to determine statistical significance with Two-stage linear step-up procedure of Benjamini, Krieger and Yekutieli to correct for FDR; mean±SEM shown. q<0.01 **, q<0.001 ***; mean±SEM shown.

Noting the enhanced binding strength of CAR.PDZ cells over a short period of time we next assessed calcium flux, the first step of NK cell activation. Utilizing live cell image analysis, we quantified single cell calcium flux levels (**Figure 2f, supplemental video 1**) and found a greater calcium burst in CAR.PDZ NK cells (**Figure 2g**) and relatively higher and sustained levels over 15 minutes in comparison to other NK cell populations (untransduced, CAR, CAR Δ). We next confirmed our findings in the DIPG007 glioma cell line and included another control: NK cells expressing CAR.PDZs in which the C-terminal amino acid of the PDZbm was mutated, to render it non-functional (CAR.PDZmut). The greatest calcium flux was observed in CAR.PDZ NK cells compared to all other groups; further, CAR Δ, CAR, and CAR.PDZmut had similar calcium flux levels (**Figure 2h**). These results directly link a functional PDZbm as being critical to enhance NK cell activation.

Given the amplified calcium influx and signaling via ZAP70 of CAR.PDZ NK cells we sought to determine the functional consequences. We first explored cytokine production upon target cell interaction and employed a 4-hour co-culture assay with A549 cells analyzing single cell secretomic profiles for each CAR construct utilizing an IsoLight™ system (**Figure 3a**). In particular, there was a notable increase in the frequency of perforin- and IFNy-secreting CAR.PDZ NK cells in comparison to CAR NK cells (**Figure 3b**). The latter is consistent with the observed amplified signaling via ZAP70 in CAR.PDZ NK cells, since IFNy production typically takes the highest level of activation stimulus and time^26^. We further observed an increased frequency of GM-CSF, TNFα, MIP-1α, and MIP-1β producing CAR.PDZ NK cells versus CAR NK cells; however, this did not reach statistical significance (**Figure 3b**). CAR.PDZ NK cells exhibited greater polyfunctionality, as judged by their ability to secrete multiple cytokines, in comparison to other NK cell populations (**Figure 3c**). To assess the quality of cytokine production, we calculated the polyfunctional strength index (PSI), which accounts for the frequency of cells secreting cytokine and the relative intensity of cytokine produced. We observed that the IFNy PSI was highest in CAR.PDZ NK cells (**Figure 3d**), which mirrored the secretion frequency data, and that the ‘effector cytokine’ group PSI was increased in CAR.PDZ versus CAR NK cells (**Figure 3e**). Finally, spectral t-SNE mapping of the signal intensity of selected cytokines (GM-CSF, IFNγ, IL-8, TNFα), chemokines (MCP-1, MIP-1α, MIP-1β) and molecules of cytolytic granules (granzyme B, perforin) revealed a distinct grouping of cells with a unique mapping of CAR.PDZ NK cells compared to other NK cell populations (**Figure 3f)**. Intriguingly, we observed that CARΔ NK cells secreted more effector molecules than untransduced NK cells and in some instances similar levels to NK cells expressing functional CARs. Increased target cell binding of NK cells by CAR Δ is the most likely explanation of our finding since it would give endogenous NK cell receptors a greater opportunity to engage cognate ligands and activate NK cells.

**Figure 3:**
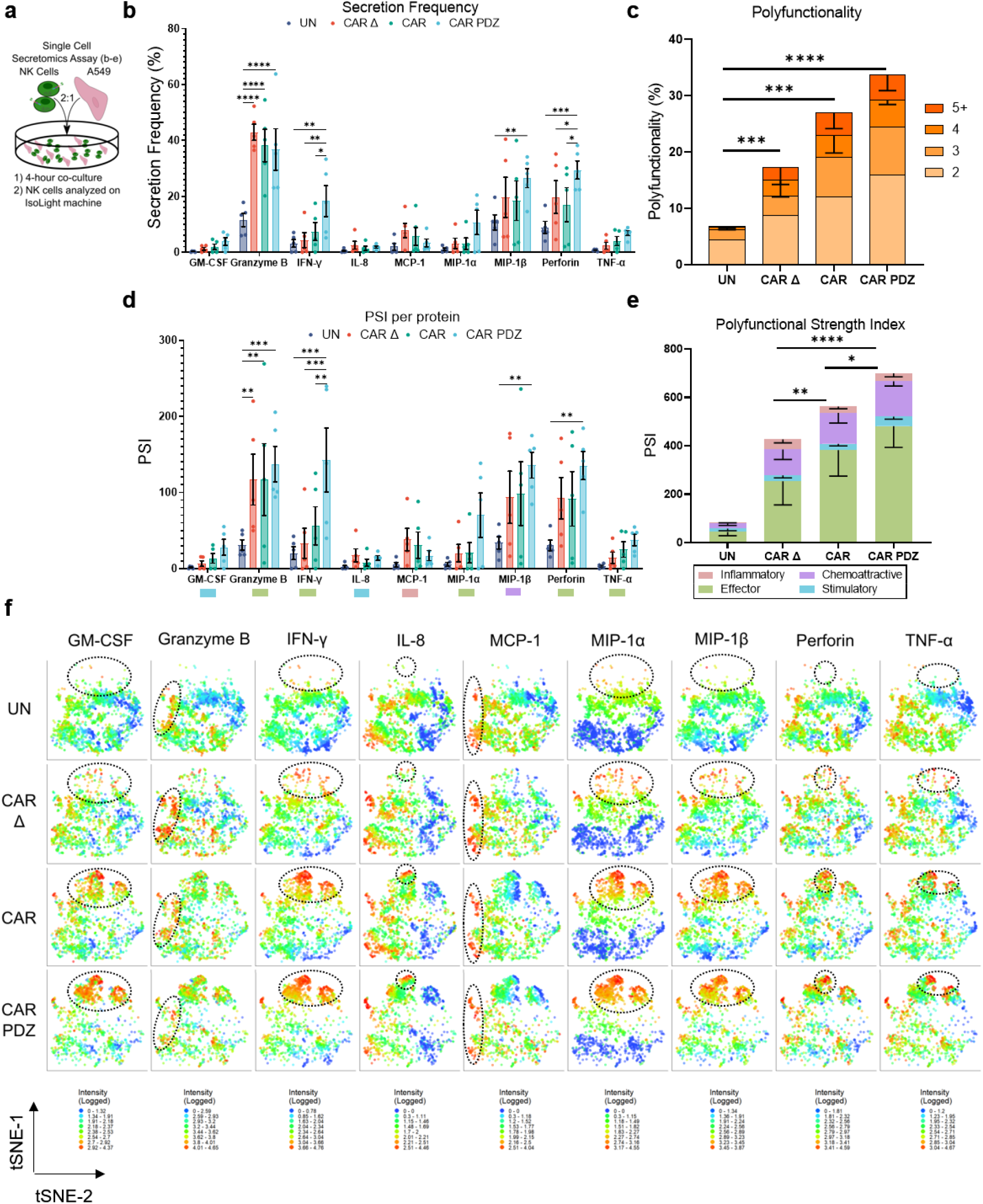
CAR.PDZ NK cells have enhanced and distinct cytokine production. (a) Schematic overview of experimental conditions for secretomics analysis. Single cell secretomic analysis using an IsoLight machine depicting the polyfunctionality of NK cells after exposure to A549 target cells for 4-hours. (b) Secretion frequency of selected cytokines from the 32 analyte IsoLight Chip. Two-Way ANOVA was used to determine statistical significance with Two-stage linear step-up procedure of Benjamini, Krieger and Yekutieli to correct for FDR. Statistical difference delineated by q<0.05 *, q<0.01 **, q<0.001 ***, q<0.0001 ****. n=5 d, mean±SEM shown. (c) All CAR constructs produced significantly more cytokines on the 2-analyte level compared to UN. Two-Way ANOVA was used to determine statistical significance with Two-stage linear step-up procedure of Benjamini, Krieger and Yekutieli to correct for FDR. Statistical difference delineated by q<0.0001 ****. n=5, mean±SEM shown. (d) Polyfunctional Strength Index (PSI) of CAR-NK cells detailing cytokine categories of NK cells from (b) individual cytokines driving PSI variance with indicated colors defining (c). Two-Way ANOVA was used to determine statistical significance with Two-stage linear step-up procedure of Benjamini, Krieger and Yekutieli to correct for FDR. Statistical difference delineated by q<0.05 *, q<0.01 **, q<0.001 ***, q<0.0001 ****. n=5, mean±SEM shown. (e) Only the effector cytokine group was significantly different between CAR.PDZ and CAR or CAR Δ NK cells. Two-Way ANOVA was used to determine statistical significance with Two-stage linear step-up procedure of Benjamini, Krieger and Yekutieli to correct for FDR. Statistical difference delineated by q<0.05 *, q<0.01 **, q<0.001 ***, q<0.0001 ****. n=5, mean±SEM shown. (f) tSNE plots of the secretomic analysis performed in Figure 2a-e. tSNE plots revealed distinct cytokine secretion profiles and patterns for each construct tested. These plots highlight the increased secretion frequency and quantity. Log transformed secretion value intensities are delineated. Dashed ellipses indicate groups of highest secretion. Color spectra vary per cytokine from 0 to the highest value in each group, n=5.

We next explored the cytolytic activity in CAR NK cells in 2D and 3D culture systems. In a standard 24 hour 2D MTS assay (**Figure 4a**), CAR.PDZ NK cells exhibited superior cytolytic activity against A549 tumor cells in comparison to CAR NK cells at all evaluated effector to target (E:T) ratios, except for the highest E:T ratio (**Figure 4b**). Assessing the mutated PDZbm CAR.PDZmut NK cells revealed a similar killing profile to standard CAR NK cells and a significantly reduced capacity compared to functional CAR.PDZ NK cells (**Figure 4c**). Conversely, we explored the functional consequences of CAR.PDZ binding to Scribble by blocking this interaction with a peptide Scribble PDZ antagonist, which resulted in reduced cytotoxicity (**Figure 4d**). We observed reduced cytolytic activity in all constructs which was expected given the ubiquitous nature of Scribble’s role in cell polarity and the high affinity peptide binding of Scribble’s PDZ domain. However, the greatest inhibitory effect was seen in CAR.PDZ NK cells as quantified by area between the curves analysis (**Figure 4d,e**).

**Figure 4:**
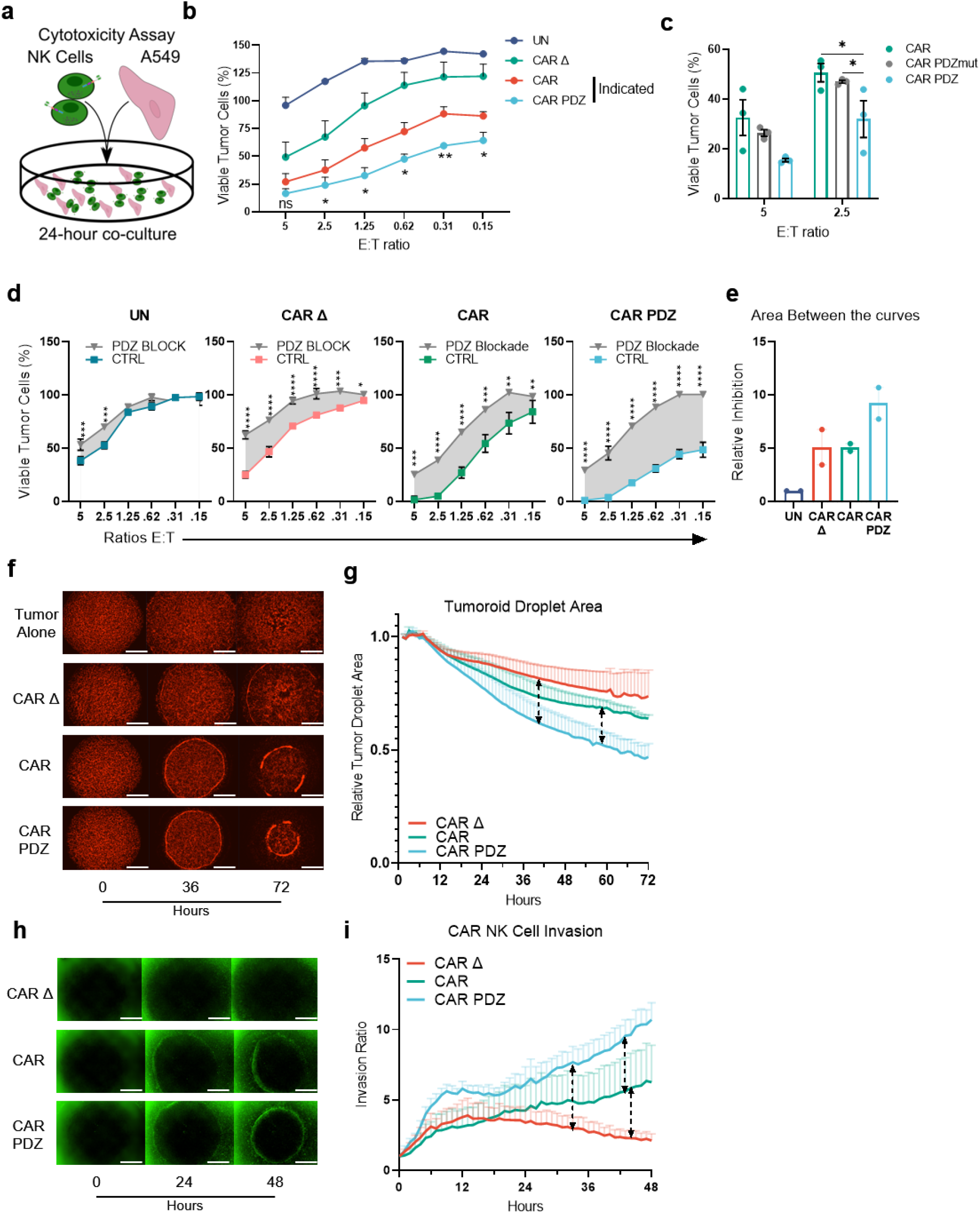
CAR.PDZ NK cells have enhanced cytolytic activity and invasive properties. (a) Cytotoxicity assay scheme with A549 lung adenocarcinoma cell viability determined by a chromogenic MTS assay after 24-hour co-culture with NK cells. (b) CAR vs CAR.PDZ NK cells: significant differences at all effector to target (E:T) ratios except for 5:1. All other comparisons are significantly different. Two-Way ANOVA was used to determine statistical significance with Two-stage linear step-up procedure of Benjamini, Krieger and Yekutieli to correct for FDR. Statistical difference delineated by q<0.05 *, q<0.01 **, q<0.001 ***, q<0.0001 ****. n=3, mean±SEM shown. (c) CAR vs CAR.PDZ vs CAR.PDZmut NK cells: significant differences at all effector to target (E:T) ratios 2.5:1. Two-Way ANOVA was used to determine statistical significance with Two-stage linear step-up procedure of Benjamini, Krieger and Yekutieli to correct for FDR. Statistical difference delineated by q<0.05 *, q<0.01 **, q<0.001 ***, q<0.0001 ****. n=3, mean±SEM shown. (d) Untransduced (UN) NK cells treated with 10µM PDZ blocking or control (CTRL) peptides. CAR Δ NK cells treated with 10µM PDZ blocking or control (CTRL) peptides. CAR NK cells treated with 10µM PDZ blocking or control (CTRL) peptides. CAR.PDZ NK cells treated with 10µM PDZ blocking or control (CTRL) peptides. Two-Way ANOVA was used to determine statistical significance with Two-stage linear step-up procedure of Benjamini, Krieger and Yekutieli to correct for FDR. Statistical difference delineated by q<0.05 *, q<0.01 **, q<0.001***, q<0.0001 ****. technical triplicates with mean±SEM shown. (e) Area between the curve analysis (shaded regions in d) for two independent experiments normalized to untransduced. (f) Representative images from a tumoroid droplet cytotoxicity assay using the Incucyte S3 imager. 143b osteosarcoma cells in mCherry were embedded in collagen and NK cells were layered around the droplet in Matrigel in a 96 well plate. Co-cultures were imaged hourly for 3 days. White bars indicate 1mm. (g) Quantification of the resulting tumor reduction as determined by tumor area normalized to the 1-hour (hr) mark. CAR.PDZ vs CAR Δ NK cells: significant differences starting at 40 hrs of culture; CAR.PDZ vs CAR NK cells: significant differences starting at 58 hrs of culture. Two-Way ANOVA was used to determine statistical significance with Two-stage linear step-up procedure of Benjamini, Krieger and Yekutieli to correct for FDR. Dashed lines indicate comparison groups and what hour they become statistically different at least q<0.05 *. n=3, mean±SEM shown. (h) Representative images from a tumoroid droplet invasion assay using the Incucyte S3 imager. 143b osteosarcoma cells in mCherry were embedded in collagen and NK cells were layered around the droplet in Matrigel labeled in green in a 96 well plate. Co-cultures were imaged hourly for 2 days. White bars indicate 1mm. (i) NK cell invasion ratio represents the counted NK cells per mm^2^ and normalized to the 1-hour mark. CAR.PDZ vs CAR Δ NK cells: significant differences starting at 33-hours of co-culture; CAR.PDZ vs CAR NK cells: significant differences starting at 43-hours of co-culture; CAR vs CAR Δ NK cells: significant differences starting at 44-hours of co-culture. Two-Way ANOVA was used to determine statistical significance with Two-stage linear step-up procedure of Benjamini, Krieger and Yekutieli to correct for FDR. Dashed lines indicate comparison groups and what hour they become statistically different at least q<0.05 *; technical triplicate with mean±SEM shown.

In a 3D assay, which consisted of a mixture of mCherry-positive 143b osteosarcoma cells and collagen in droplets surrounded by a ring of NK cells in Matrigel (**Figure 4f**), CAR.PDZ NK cells reduced tumoroid size to a greater degree than CAR NK cells (**Figure 4g**). NK cells were then labeled with a green, fluorescent dye and tracked over 48 hours. CAR.PDZ cells also exhibited an enhanced ability to migrate to the droplet and invade to the center of the well in comparison to CAR NK cells (**Figure 4h,i; supplemental videos 2-4**). Thus, adding a PDZbm to CARs enhances not only their cytolytic potential, but also their migratory activity consistent with the known biology of Scribble in cell polarity and migration^6^.

Finally, we compared the anti-tumor efficacy of untransduced, CAR Δ, CAR, and CAR.PDZ NK cells targeting EphA2 in two solid tumor xenograft models. We started with an A549 lung adenocarcinoma model in which tumor cells were implanted subcutaneously and treated with NK cells intravenously 14 days later (**Figure 5a**). Untreated tumors and tumors treated with untransduced NK cells showed rapid outgrowth. In contrast, CAR Δ, CAR, and CAR.PDZ NK cells had anti-tumor activity, including complete responses (CRs) for CAR and CAR.PDZ NK cells (**Figure 5b**). Only mice treated with CAR.PDZ NK cells had an improved overall survival benefit in comparison to CAR Δ NK cell treated mice (**Figure 5c**). While the median survival of CAR.PDZ NK cell treated mice was 16 days greater than for CAR NK cell treated mice, this did not reach statistical significance (**Figure 5c**). We next evaluated the same CAR NK cell populations in a locoregional osteosarcoma model. Mice were injected intraperitoneally with firefly luciferase expressing LM7 cells followed by injection of NK cells on day 7 (**Figure 5d**). Only, CAR.PDZ and CAR NK cells induced tumor regression, including CRs, as judged by bioluminescence imaging (**Figure 5e**). Mice treated with CAR.PDZ NK cells had a distinct survival advantage in comparison to all other treatment groups including CAR NK cells (**Figure 5f**). We extended these findings to the 143b osteosarcoma model using CARs and CAR.PDZs specific for EphA2 or B7-H3 (**Figure 5g**). Post infusion of untransduced, CAR Δ, CAR, or CAR.PDZ NK cells tumors continued to grow rapidly except in mice that had received CAR.PDZ NK cells (**Figure 5h**), resulting in a significant survival advantage (**Figure5i,j**).

**Figure 5:**
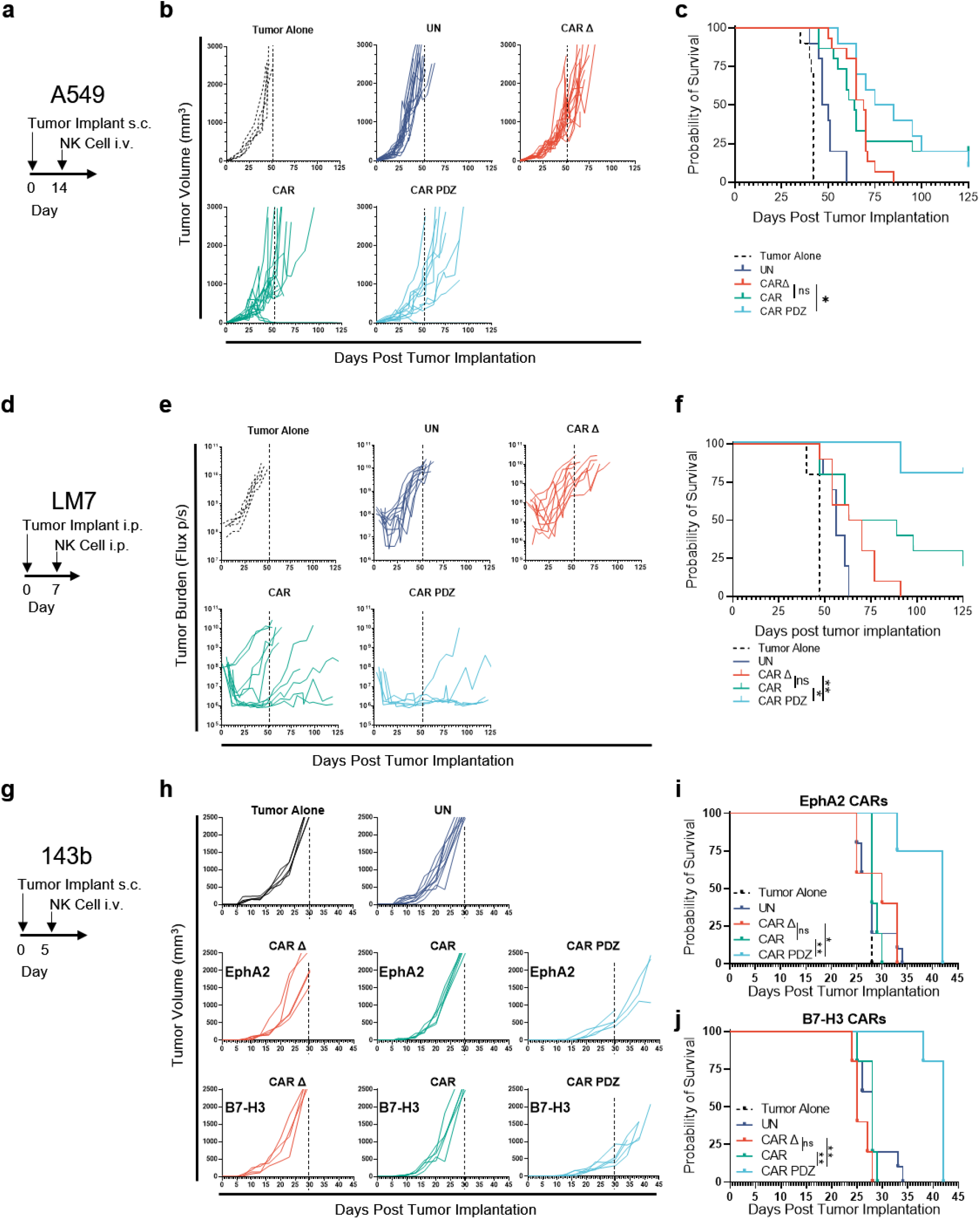
CAR.PDZ NK cells extend survival and eradicate solid tumors *in vivo*. (a) A549 model timeline. (b) 2×10^6^ A549 tumor cells were mixed in pure Matrigel and injected subcutaneously (s.c.) into the dorsal flank of male and female NSG mice. 14 days later mice were treated with a single 10×10^6^ intravenous (i.v.) injection of NK cells. Each group of mice is n=5-15 merged from two-three independent experiments. (c) Kaplan-Meier curves of mice from (a) median survival rates for each group (days): Tumor Alone: 42, UN: 48.5, CAR Δ: 69, CAR: 64, CAR.PDZ: 80. Log-rank test was used to determine significance. Statistical difference delineated by p<0.05 *. (d) LM7 locoregional model timeline. (e) 1×10^6^ LM7.ffluc tumor cells were injected intraperitoneally (i.p.) into male and female NSG mice. 7 days later mice were treated with a single 10×10^6^ i.p. dose of NK cells. Tumors were measured by bioluminescence imaging with total photon flux depicted. Each group of mice is n=5-10 merged from two independent experiments. (f) Kaplan-Meier curves of mice from (b) median survival rates for each group (days): Tumor Alone: 47, UN: 56, CAR Δ: 66.5, CAR: 76, CAR.PDZ: undefined. Log-rank test was used to determine significance. Statistical difference delineated by p<0.05 *, <0.01 **. (g) 143b model timeline. (h) 1×10^6^ 143b tumor cells were mixed in pure Matrigel and injected subcutaneously (s.c.) into the dorsal flank of male and female NSG mice. 5 days later mice were treated with a single 10×10^6^ intravenous (i.v.) injection of NK cells. Each group of mice is n=4-10 Tumor alone and Untransduced groups are shared between CAR target groups. (i) Kaplan-Meier curves of EphA2 targeted CAR NK treated mice from (h) median survival rates for each group (days): Tumor Alone: 28, UN: 28, CAR Δ:30, CAR: 28, CAR.PDZ: 4.2 Log-rank test was used to determine significance. Statistical difference delineated by p<0.05 *, <0.01 **. (j) Kaplan-Meier curves of B7-H3 targeted CAR NK treated mice from (h) median survival rates for each group (days): Tumor Alone: 28, UN: 28, CAR Δ:25, CAR: 28, CAR.PDZ: 4.2 Log-rank test was used to determine significance. Statistical difference delineated by p<0.01 **.

We further evaluated the functional persistence of CAR NK cell in mice that had achieved a CR in both models. In the A549 model, mice were re-challenged with the original tumor cell dose ∼4 months after initial therapy. Palpable tumor nodules were measured 7 days post injection; however, large tumors never formed, and nodules had disappeared by day 14 (**Extended Data Figure 6a**). Likewise, LM7 tumors were rejected post rechallenge ∼4 months after initial therapy (**Extended Data Figure 6b,c**). Thus, CAR NK cells persisted and retained anti-tumor capabilities even though NSG mice do not endogenously produce cytokines that support NK cell survival.

Persistence has been long thought as a hurdle of NK cell therapies, and many investigators are working on deciphering and attempting to solve this problem^27^, including engineering NK cells to express IL-15, priming NK cells with cytokines to induce a “memory-like” phenotype, or culturing NK cells with various ligand expressing feeder cells^28^ to enhance persistence. Our culturing system involves low dose interleukin (IL)-2, 10IU/mL, and a single, initial feeder cell stimulation. Utilizing a low stimulation level, putatively, reduces the potential for dependence on high levels of cytokine to maintain growth and survival post adoptive transfer into a harsh environment.

NK cells have been derived from peripheral blood (PB)^29^, cord blood (CB)^30^, induced pluripotent stem cell (iPSC)^31^, or existing cell lines, e.g. NK92^32^. NK cells derived from these sources have been evaluated in clinical studies with an encouraging safety record to date^28,33^. We focused in our study on PB-derived CAR NK cells since they are readily available from healthy donors. PB has been largely overlooked as a CAR NK cell source, especially in the context of solid tumor-redirected CAR NK cell therapy^34^. For example, there are only three preclinical publications that report *in vivo* experiments, all of which utilized repeat dosing regimens, high dose cytokine supplementation, or both^34^. Thus, our encouraging results should provide impetus for the active exploration of PB-derived CAR NK cells. NK cells, regardless of source, need migratory, polarization, and internal scaffolding programs to be effectively employed as anti-cancer therapeutics.

We selected to exploit the naturally occurring cell polarity requirements for immune cell recognition of target cells^35^. Cell polarity is a tightly regulated process that has limited targets to augment without unintended consequences^36^. To this end, we found that the extant PDZbm of CRTAM would be an ideal candidate to explore for our new synapse tuned design. Further, CRTAM is considered to be a late phase polarity protein that aids in cell signaling after antigen recognition^4^. Thus, we posited that introducing the PDZbm domain of CRTAM into CARs would create a more efficient signaling CAR and imbue effector cells with enhanced functionality and longevity. Thus, our developed CAR.PDZ should enhance the effector function of all NK cell products. Finally, since PDZbms and their interactions with scaffolding proteins is evolutionarily conserved, it is likely that PDZbms will enhance the effector function of other CAR-expressing immune cells, including T cells, that are actively being explored as immunotherapeutics. Noting our observed increases in avidity and IFNγ secretion, we believe that our PDZbm domain has the potential to also increase the efficacy of CAR T cells for solid tumors given a recent publication^37^ that detailed that both factors are critically important for their antitumor efficacy. Additionally, adding this domain to the C-terminal of other endogenous receptors, such as CD16, may enhance antibody-based therapeutics through antibody dependent cellular cytotoxicity.

Exploration into CAR synapse tuning is at present a sparsely populated field with studies observing differences between various standard CAR designs such as bi-specific CARs^38,39^ and CARs with different costimulatory domains^40,41^. We contend that our report is the first synapse directed proof-of-concept study that reveals distinct advantages by adding a domain designed to augment the synapse. A plethora of research has been performed in optimizing CAR designs looking at external factors such as single chain variable fragment (scFv) affinity and which costimulatory domain compliments the overall activation domain. However, these efforts have been in the context of an inefficient synapse. Orienting our focus to the inside of the cell to determine optimal organization of the synapse takes an orthogonal and complimentary approach to current research efforts. Re-evaluating previously discarded CAR designs with a new lens of synapse tuning may give life to previously shelved ideas. In conclusion, we demonstrate here that an optimal CAR endodomain might not only have to include costimulatory and activation domains, but also a PDZbm-based ‘anchoring domain’. Thus, synapse tuning via anchor domains represents a new era in CAR design and a fertile realm to explore for CAR-based immunotherapies.

## Acknowledgements

The work was supported by the St. Jude Sumara Fellowship (PC), the American Lebanese Syrian Associated Charites (SG), ChadTough Defeat DIPG Foundation (GK), (NINDS) Grant R01NS121249 (GK), the Rally Foundation for Childhood Cancer Research (LJT), The Garwood PostDoctoral Fellowship (JIV) and the National Institutes of Health (NIH)/National Cancer Institute (NCI) grant P30 CA021765. Animal imaging was performed by the Center for In Vivo Imaging and Therapeutics, which is supported in part by NIH grants P01CA096832 and R50CA211481. Cellular images were acquired at SJCRH Cell & Tissue Imaging Center which is supported by St. Jude and NCI P30 CA021765. Gene editing of cell lines was performed by the Center for Advanced Genome Engineering (CAGE), which is supported in part by NCI P30 CA021765. The content is solely the responsibility of the authors and does not necessarily represent the official views of the NIH.

## Author Contributions

P.J.C. conceptualization, performed experiments, analyzed data, and wrote manuscript. J.I. performed confocal microscopy and analysis. G.K. provided DIPG007 model setup for immune cell studies and acquired funding, L.J.T. developed and performed Halo assay. S.G. acquired funding, supervised, and wrote manuscript.

## Conflict of Interest

S.G. and P.J.C. have patent applications in the fields of NK and T-cell and/or gene therapy for cancer. S.G. has a research collaboration with TESSA Therapeutics, is a DSMB member of Immatics, and was on the scientific advisory board of Tidal.

## Corresponding Author

Correspondence should be addressed to Peter J Chockley

## Methods

### Cell Lines

143b osteosarcoma and A549 lung cancer cell lines were obtained and grown as per American Type Culture Collection (ATCC, Manassas, VA, USA) instructions. LM7, a metastatic osteosarcoma cell line, was provided by Dr. Eugenie Kleinerman (MD Anderson Cancer Center, Houston, Texas, USA) in 2011. The generation of LM7 cells expressing an enhanced green fluorescent protein firefly luciferase fusion protein (LM7.eGFP.ffLuc) was previously described^42^. K562 with modified membrane bound IL-15 and 4-1BB ligand^43^, feeder cells, were a generous gift from Dr. Dario Campana (National University of Singapore), and grown in IMDM media with 10% fetal bovine serum (FBS; Hyclone Laboratories, Chicago, IL, USA). The EphA2 KO A549 cell line was generated with CRISPR/Cas9 technology using a published method^44^. HSJD-DIPG007 (DIPG007) cells were cultured as described previously^45^. Cell lines were validated with short tandem repeat profiling performed by ATCC.

### Generation of Retroviral Vectors

The generation of the SFG retroviral vectors encoding EphA2-CARs with a CD28 costimulatory domain (CAR), a nonfunctional EphA2-CAR without a signaling domain (CAR Δ) were previously described^18^. B7-H3 CARs were generated similar and previously described^24^. In-Fusion cloning (Takara Bio, Kusatsu, Shiga, Japan) was used to generate the CAR.PDZ with a PDZbm attached to the C-terminus of CD3ζ domain and site directed mutagenesis Q5(New England Biolabs, Ipswich, MA, USA) was used to do a point mutation to exchange the final Valine to an Alanine to create the CAR.PDZmut construct.

CAR.PDZ additional sequence containing PDZbm:

HPMRCMNYITKLYSEAKTKRKENVQHSKLEEKHIQVP<u>ESIV*</u>

CAR.PDZ additional sequence containing mutated PDZbm:

HPMRCMNYITKLYSEAKTKRKENVQHSKLEEKHIQVP<u>ESIA*</u>

The sequence of the final construct was verified by Sanger sequencing (Hartwell Center, St. Jude Children’s Research Hospital). Retroviral particles were generated as previously described^46^ by transient transfection of HEK293T cells (ATCC) with the EphA2-CAR encoding SFG retroviral vectors, Peg-Pam-e plasmid encoding MoMLV gag-pol, and a plasmid encoding the RD114 envelope protein. Supernatants were collected after 48 hours, filtered, and snap-frozen for later transduction of NK cells.

### NK cell Activation, Expansion, and Genetic Modification

Human peripheral blood mononuclear cells (PBMCs) were obtained from whole blood of healthy donors under an IRB approved protocol at St. Jude Children’s Research Hospital (St. Jude), after informed consent was obtained in accordance with the Declaration of Helsinki, or from de-identified elutriation chambers of leukapheresis products obtained from St. Jude donor center. Donors were less than haplo-identically matched by HLA typing. Cells were subjected to ACK Red Blood cell lysis and Ficoll Hypaque (Sigma-Aldrich, St. Louis, MO, USA) gradient separation. Cellular subtype analysis was performed with BD whole blood analysis kit on a BD Lyric flow cytometer (Becton-Dickinson, Franklin Lakes, NJ, USA). PBMCs were depleted of CD4 and CD8 using standard MACs magnetic beads (CD4: 130-045-101, CD8: 130-045-201, Miltenyi Biotec, Bergisch Gladbach, North Rhine-Westphalia, Germany). Cells were aliquoted in freezing media with 10% DMSO at 1×10^7^ cells per mL and stored in liquid nitrogen vapor phase until use. 150 Gray cesium-irradiated feeder cells were added to thawed CD4/8-depleted PBMCs at a ratio of 5-10:1 feeder to NK cells. Cells were grown in Stemcell Genix (20802-0500, Cellgenix, Portsmouth, MA, USA) growth media with 20% FBS and 10 units/mL of IL-2, (Peprotech, Rocky Hill, NJ, USA) (complete growth media). After 5-7 days cells were phenotyped and used for downstream experiments.

Genetically modified NK cells were generated as follows: Supernatants containing retroviral particles encoding CAR constructs were spun at 2000g in retronectin (T100B, Takara Bio) coated non-tissue culture 24-well plates for 90 minutes. Supernatants were removed and 250,000 NK cells were seeded per well in a volume of 1 mL of complete growth media. NK cells were incubated for 24 hours and then removed and cultured complete growth media. Modified NK cells were expanded in G-Rex 6 well plates for 10-14 days (#80240MWilson Wolf, New Brighton, MN, USA). NK cell transgene expression was assessed 3-7 days post-transduction.

### Flow Cytometry

250,000 NK cells were collected and washed twice in DPBS. Surface EphA2 or B7-H3-CAR detection was determined via immunolabeling with anti-F(ab’)2-AF647 (109-606-006, Jackson Labs, Bar Harbor, ME, USA; 1:100), was utilized for detection on a BD FACS Lyric machine and analyzed with FlowJo v10 (BD). Immunophenotyping antibodies are listed in extended table 1.

### Single Cell Secretomics Assay

Briefly, 100,000 NK cells were labeled with cell trace violet 1:500 (ThermoFisher Scientific, Waltham, MA, USA), and co-cultured at a 2:1 ratio with A549 targets for 4-hours in a 48 well plate. We found that 24-hour co-cultures resulted in increased tumor cell death and reduced cytokine detection. NK cells were removed and washed twice with PBS and resuspended in complete growth media without IL-2. These cells were loaded onto a IsoCode single cell secretomic chip and run on the IsoLight machine that detects 32 distinct proteins^47^. Results were analyzed on IsoSpeak version 2.8.1.0 (IsoPlexis, Branford, CT, USA).

### Cytotoxicity Assays

NK cells were cytokine starved for 24-hours prior to co-culture with target cells to reduce non-specific killing of target cells. 3,000 target (A549) cells were cocultured with effectors at indicated effector to target (E:T) ratios for 24-hours in a 96 well plate. Cytotoxicity was quantified by a chromogenic MTS assay measured on a plate reader (Tecan, Männedorf, Switzerland) detecting remaining viable adherent tumor cells.

### Peptide Blockade

Peptides were synthesized by the Macromolecular Synthesis Core (Harwell Center, St. Jude) purity was 97% and 95% via HPLC, respectively:

Negative Control Sequence: NH_2_-<u>RQIKIWFQNRRMKWKK</u>RSWFEAWA-COOH

Scribble PDZ Blocking Sequence: NH_2_-<u>RQIKIWFQNRRMKWKK</u>RSWFETWV-COOH

Underlined portions of sequences are from the Antennapedia protein which has been shown to allow peptide shuttling into cells and effectively block CRTAM binding^48^. NK cells were treated with 10micromolar of peptides for 24-hours prior to co-culture.

### Confocal Microscopy

#### Ag-coated coverslip preparation and NK activation

Antigen coated coverslips were prepared using N1 coverslips (Fischer Scientific: #12-545-80P), which were coated with 0.5 µg/mL of rhEphA2 (R&D Systems, Minneapolis, MN, USA: #3035-A2-100) or poly-L-lysine (Sigma: #P4707) overnight at 4°C. Then, they were washed with PBS and filled with media until NK cell seeding. 200,000 NK cells were plated onto the precoated coverslips at different time points in a cell culture incubator (37°C/5%CO_2_). After activation, NK cells were washed with cold PBS and fixed with 4% paraformaldehyde (PFA, Electron microscopy sciences #15710) for 10 minutes at room temperature. Fixed cells were washed twice with PBS and the remaining PFA was inactivated with blocking buffer (PBS-2% BSA (Sigma: #A9418), 1.5M Glycine (Sigma: #G8898)) for 10 min at room temperature. Cells were permeabilized by adding permeabilization buffer (PBS, 0.2%BSA, 0.05%Saponin; Sigma: #47036) for 20 minutes at room temperature. Cells were washed twice with permeabilization buffer prior primary antibody incubation diluted in permeabilization buffer, following manufacturer instructions. All the primary antibodies were incubated at 4°C overnight. Cells were washed with permeabilization buffer and incubated with secondary antibodies for 2-hours at room temperature. Finally, cells were washed with permeabilization buffer and PBS before to let them dry for 1-hour at room temperature. Then, coverslips were mounted onto slides using fluoromount (Thermofisher Scientific: #00-4958-02).

#### Antibodies and probes

Primary antibodies and probes with their dilutions: Anti-human Lamp1 (1:50) (Abcam: #ab25630), Anti-Human pZAP70 (1:50) (Cell Signaling Technology: #2701L), (Abcam: Ab6160), Phalloidin-AlexaFluor647 (1:200) (Thermofisher Scientific: #A22287), Anti-Human CD3e-AlexaFluor 647 (1:100) (Biolegend: #100209), Anti-Human Scribble (1:100) (Cell Signaling Technology, Danvers, MA, USA: #4475), Anti-WASp (1:100) (AbCam: #ab75830). Secondary antibodies with their dilutions: Anti-Rabbit AlexaFluor 488 (1:200) (Thermofisher Scientific, #A32731), Anti-Mouse AlexaFluor 568 (1:200) (Thermofisher Scientific, #A-11004).

#### Image acquisition and analysis

Images were acquired in a spinning disc confocal microscope (Zeiss Axio Observer with CSU-X spinning disc), and the processing and analysis was performed with FIJI (ImageJ) software^49^. Single cell images shown in the figures were cropped from larger field. Image brightness and contrast was manually adjusted. To analyze lysosome and pZAP distribution in the immune synapse, cell borders were automatically delimited by using WEKA^50^. To segmentate every single cell using F-actin signal as a template (CellTemp), an ellipse was automatically determined (CenterTemp) at the center of the CellTemp, which had a third of the CellTemp area. The recruitment at the center of the IS was calculated by dividing the fluorescence normalized by its area from CenterTemp and CellTemp, subtracting 1. Therefore, positive values mean that the fluorescence is enriched at the center, whereas negative values mean peripheral enrichment. The code is available in GitHub (https://github.com/Jorge-Ibanez-StJude/AutomatedImageAnalysis.git).

#### Live cell Calcium imaging

150,000 tumor cells were seeded onto μ-slide 8 well chambers (ibidi, Gräfelfing, Germany) (ibidi#80807) and incubated overnight at 37°C and 5% CO_2_. Tumor cells were labeled with CellBrite® Green membrane dye (Biotium Inc, Fremont, CA, USA) (1:2000) (biotum#30021) for 30 min and then washed and maintained in RPMI until image acquisition. 2×10^6^ CAR NK cells were resuspended in 1mL of PBS and labeled with CAL520 (1:500) (ATTbioquest#21130) and celltrace violet (1:1000) (Thermofisher #C34557) for 1 hour and then washed and maintained in RPMI until image acquisition.

3×10^5^ CAR NK cells were added to each well preloaded with tumor cells, and the image acquisition was initiated upon NK cells detection in the visual field. Images were acquired in a spinning disc confocal microscope (Zeiss Axio Observer with CSU-X spinning disc), using a 63X objective. The acquisition parameters were a 4D image (60 min of acquisition with 1 min of frame, and 20µm of height with a Z-step of 1µm).

The processing and analysis were performed with FIJI (ImageJ) software. Cell tracking and calcium influx were performed by using Trackmate plugin^51^ with WEKA segmentation. All tumor and CAR NK cell interaction were recorded, and calcium influx was measured as the maximum fluorescence emitted by CAL520 signal, and it was normalized by its value before the first peak of calcium influx upon tumor interaction.

### Single Cell Avidity Assay

Briefly, A549 or LM7 cells were seeded into a poly-L-lysine (Sigma) coated piezo chip from Lumicks. A549 and LM7 cells adhered for 2-hours. NK cells were labelled with celltrace FarRed (ThermoFisher) at 1:1000 dilution. The A549 laden piezo chip was loaded onto the z-MOVI single cell avidity analyzer. Labelled NK cells were injected into the chip and allowed to incubate on the A549 or LM7 cells for 5 minutes. After this time, NK cells were subjected to increasing acoustic force ramp from 0 to 1000pN over 2 minutes and 30 seconds. Individual cells were observed and the exact force requirement for detachment was recorded based on the individual cell leaving the focal plane.

### Halo Tumor Invasion Assay

Briefly, NK cells were stained with CellBrite® Green membrane dye (Biotium Inc) according to manufacturer’s instructions. They were then resuspended in a 4:3 vol:vol mixture of reduced-growth factor Matrigel (Corning, Glendale, AZ, USA) and complete RPMI without cytokines (halo matrix) at a concentration of 2×10^5^ effector cells per 5µL halo matrix. The halo matrix and resuspended cells were plated manually in a ring around the periphery of a 96-well tissue-culture treated plate (Corning) Next, 1.5% rat-tail collagen I was prepared from 3% stock (ThermoFisher Scientific), 1N NaOH, 10X PBS, and complete RPMI. 143B cells expressing mCherry were then resuspended in 1.5% collagen at a concentration of 1×10^5^ cells/1µL 1.5% collagen. An E3X Repeater® pipette (Eppendorf) with a 0.1mL Combitip advanced pipette tip was used to dispense 1µL of resuspended 143B cells in a droplet in the center of each well. After collagen and gel solidification, complete media was then layered on top for imaging with an Incucyte S3 live-cell analysis system (Sartorius, Göttingen, Germany). Imaging was performed every hour at 4X magnification using red, green, and bright field channels. For analysis, homing of effector cells was defined as the total green area in µm^2^ per image; the central tumoroid control was defined as percent total red area (µm^2^) per image normalized to hour 1 post initiation of scanning.

### In Vivo Tumor Models

Animal experiments followed a protocol approved by the St. Jude Institutional Animal Care and Use Committee. All experiments used 8- to 9-week female or male NSG mice obtained from the St. Jude NSG colony. Mice were euthanized when they reached our tumor burden limit or when they met physical euthanasia criteria (significant weight loss, signs of distress) or when recommended by St. Jude veterinary staff.

#### Sub-cutaneous tumor models

A549 cells were injected subcutaneously at 2×10^6^ cells per 100µL of Matrigel into the dorsal flanks of NSG mice. 10×10^6^ NK cells per mouse were injected intravenously on day 14. Tumor volume was calculated with the modified ellipsoidal equation (LxWxW)/2 every 5 days and mice were euthanized upon reaching a tumor volume limit of 3000mm^3^ or for humane reasons determined by St Jude veterinarians. 143b cells were injected subcutaneously at 1×10^6^ cells per 100µL of Matrigel into the dorsal flanks of NSG mice. 10×10^6^ NK cells per mouse were injected intravenously on day 5. Tumor volume was calculated with the modified ellipsoidal equation (LxWxW)/2 every 5 days and mice were euthanized upon reaching a tumor volume limit of 2500mm^3^ or for humane reasons determined by St. Jude veterinarians.

#### Locoregional LM7 osteosarcoma model

1×10^6^ LM7.eGFP.ffLuc expressing cells were injected intraperitoneally (i.p.) into the peritoneal cavity of NSG mice. 10×10^6^ NK cells per mouse were injected intraperitoneally on day 7. Mice were then imaged weekly. For imaging, mice were injected i.p. with 150 mg/kg of D-luciferin 5-10 minutes before imaging, anesthetized with isoflurane, and imaged with a Xenogen IVIS-200 imaging system (PerkinElmer, Waltham, MA, USA). The photons emitted from the luciferase-expressing tumor cells were quantified using Living Image software (Caliper Life Sciences). Total emitted photon flux (photons per second: p/s) was used to determine tumor burden and mice were euthanized upon reaching 1×10^10^ p/s.

### Used Software

IsoSpeak v2.8.1.0, GraphPad Prism v9, FlowJo v10, Fiji, Incucyte 2020A, Living Image, Oceon 1.2.1

### Statistical Analysis

Statistical analysis was performed using Graphpad Prism v9.2.0. Comparisons between two groups were determined by unpaired, two-tailed, Student’s t-Test. Three or more groups comparisons were performed with One-Way ANOVA or Two-Way ANOVA with Two-stage linear step-up procedure of Benjamini, Krieger and Yekutieli to correct for false discovery rates (FDR). This method includes the p value variance to help control for false positives and determine true discoveries based on a q value threshold <0.05. Tests are indicated in each figure legend and what value is being shown.

**Extended Data Figure 1:**
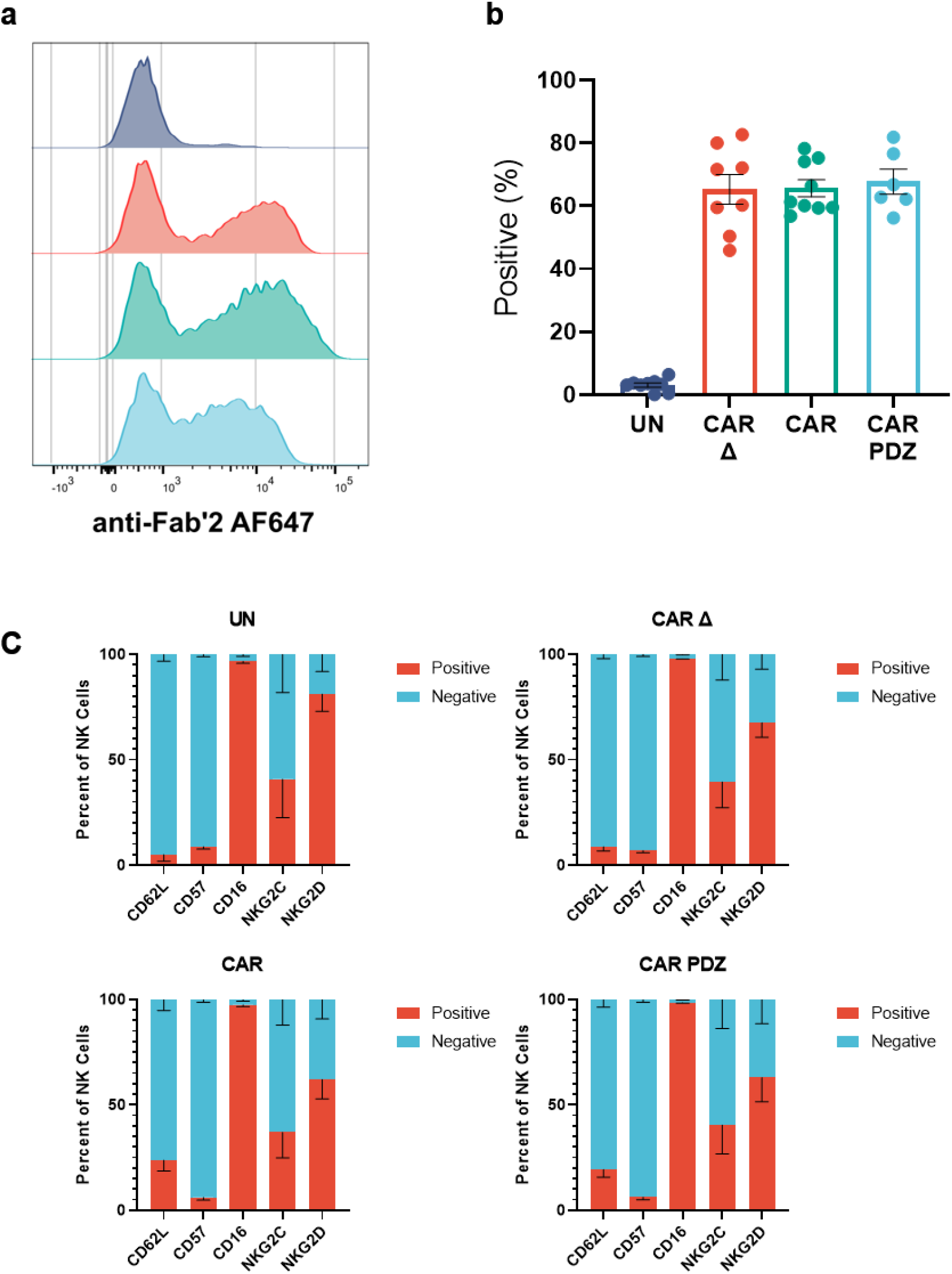
Primary NK cell transduction efficiency. (a) Representative flow cytometry histogram plots detailing surface CAR expression. (b) Quantified flow cytometry data showing percent CAR positive NK cells. n=6-9, mean±SEM shown. (c) Immunophenotype of NK cells via flow cytometry. n=4, mean±SEM shown.

**Extended Data Figure 2:**
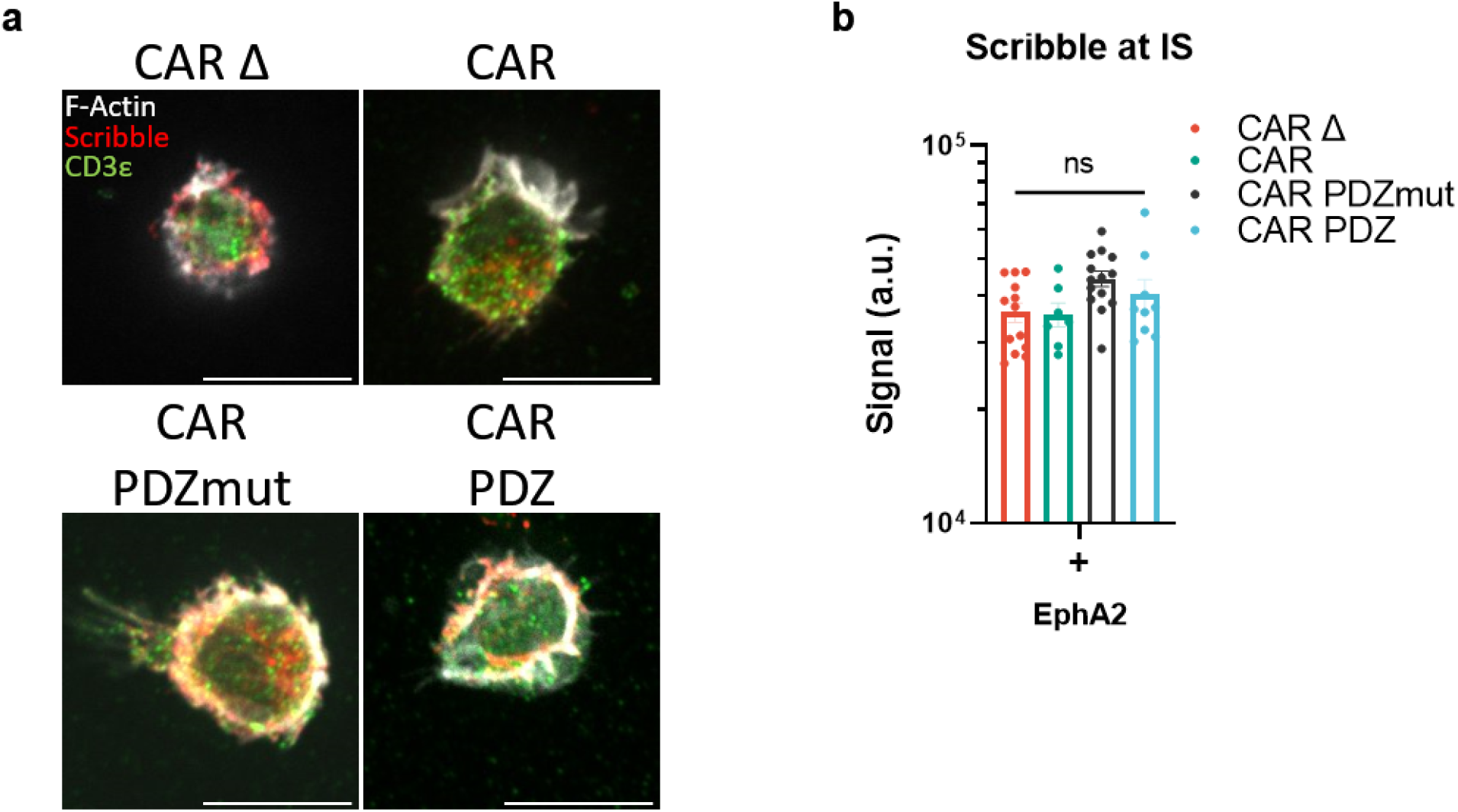
Scribble Polarization at 30 minutes. (a) Confocal images as prepared in (Figure 1) incubated for 30minutes with NK cells in various groups quantified in (b). White bars indicate 10microns. Immunolabelling of Scribble in red, CD3ε in green, and filamentous actin (F-actin) in white. (b) Scribble polarization and accumulation at the immune synapse (IS). CAR Δ; n=13, CAR; n=7, CAR.PDZmut: n=14, CAR.PDZ n=9, One-Way ANOVA was used to determine statistical significance with Two-stage linear step-up procedure of Benjamini, Krieger, and Yekutieli to correct for FDR. mean±SEM shown.

**Extended Data Figure 3:**
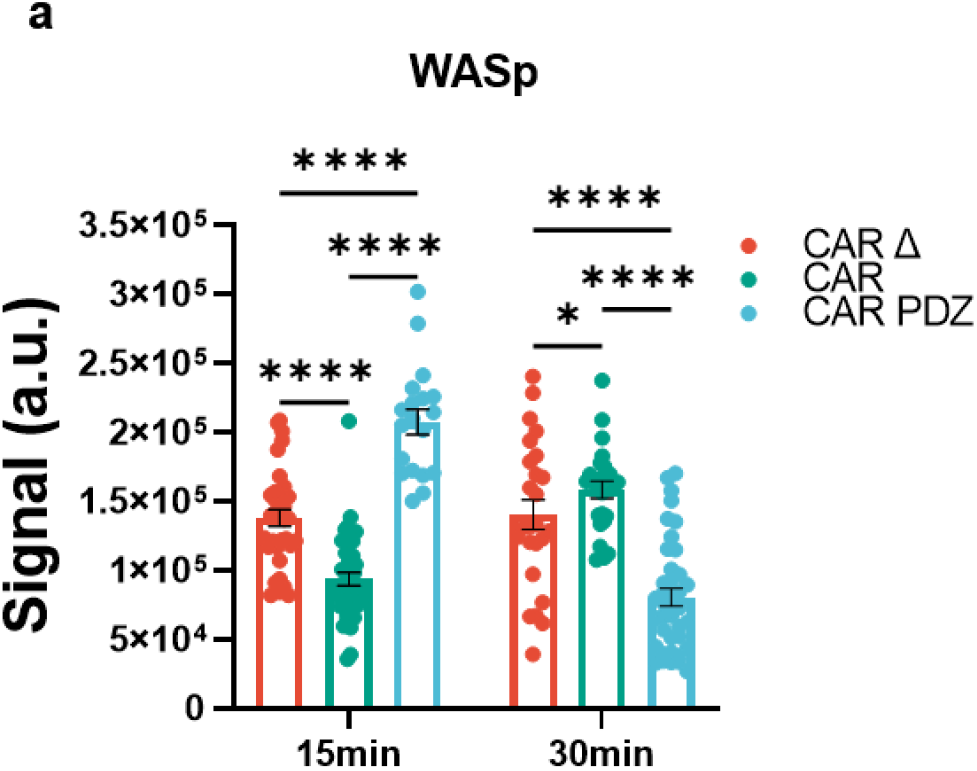
WASp Polarization at 15 and 30 minutes. (a) Confocal images as prepared in (Figure 1) incubated for 15 and 30 minutes with NK cells in various groups. Quantified WASp polarization and accumulation at the immune synapse (IS). CAR Δ; n=34 and 25, CAR; n=42 and 25, CAR.PDZ n=19 and 39, for 15 and 30 minutes, respectively. Two-Way ANOVA was used to determine statistical significance with Two-stage linear step-up procedure of Benjamini, Krieger, and Yekutieli to correct for FDR. mean±SEM shown. Statistical difference delineated by q<0.01 *, q<0.0001 ****; mean±SEM shown.

**Extended Data Figure 4:**
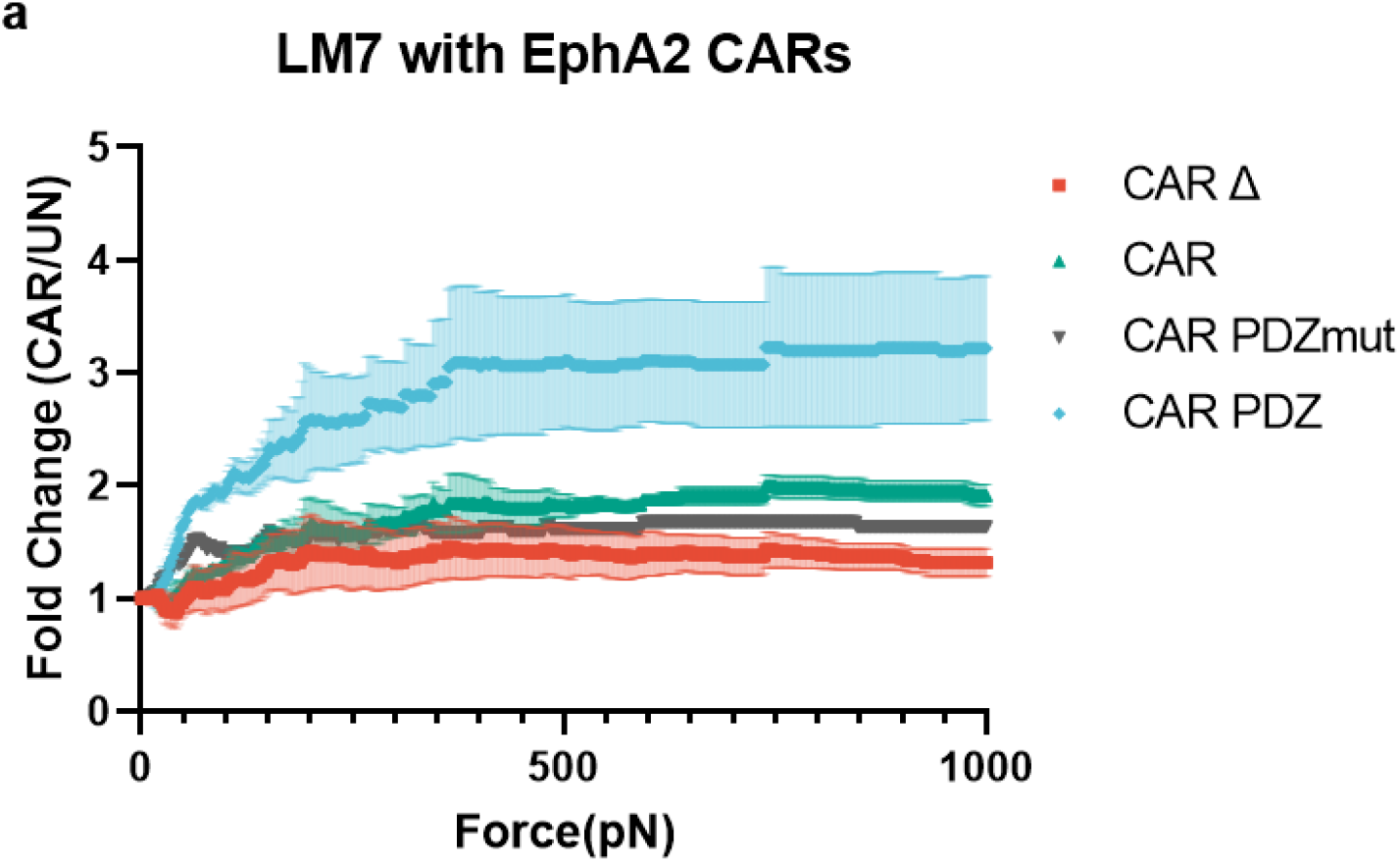
LM7 Avidity assessment with EphA2 targeting CARs. (a) Normalized fold change of CAR NK cell binding compared to untransduced NK cells. Arrowed lines indicate the point of statistical difference at CAR.PDZ vs. CAR Δ at 268pN, CAR.PDZ vs CAR at 343pN, CAR.PDZ vs CAR.PDZmut at 363pN which continued to 1000pN. The only exception to this significance was from 650 to 738pN for CAR.PDZ vs CAR.PDZmut. Two-Way ANOVA was used to determine statistical significance with Two-stage linear step-up procedure of Benjamini, Krieger and Yekutieli to correct for FDR; n=1-3, mean±SEM shown.

**Extended Data Figure 5:**
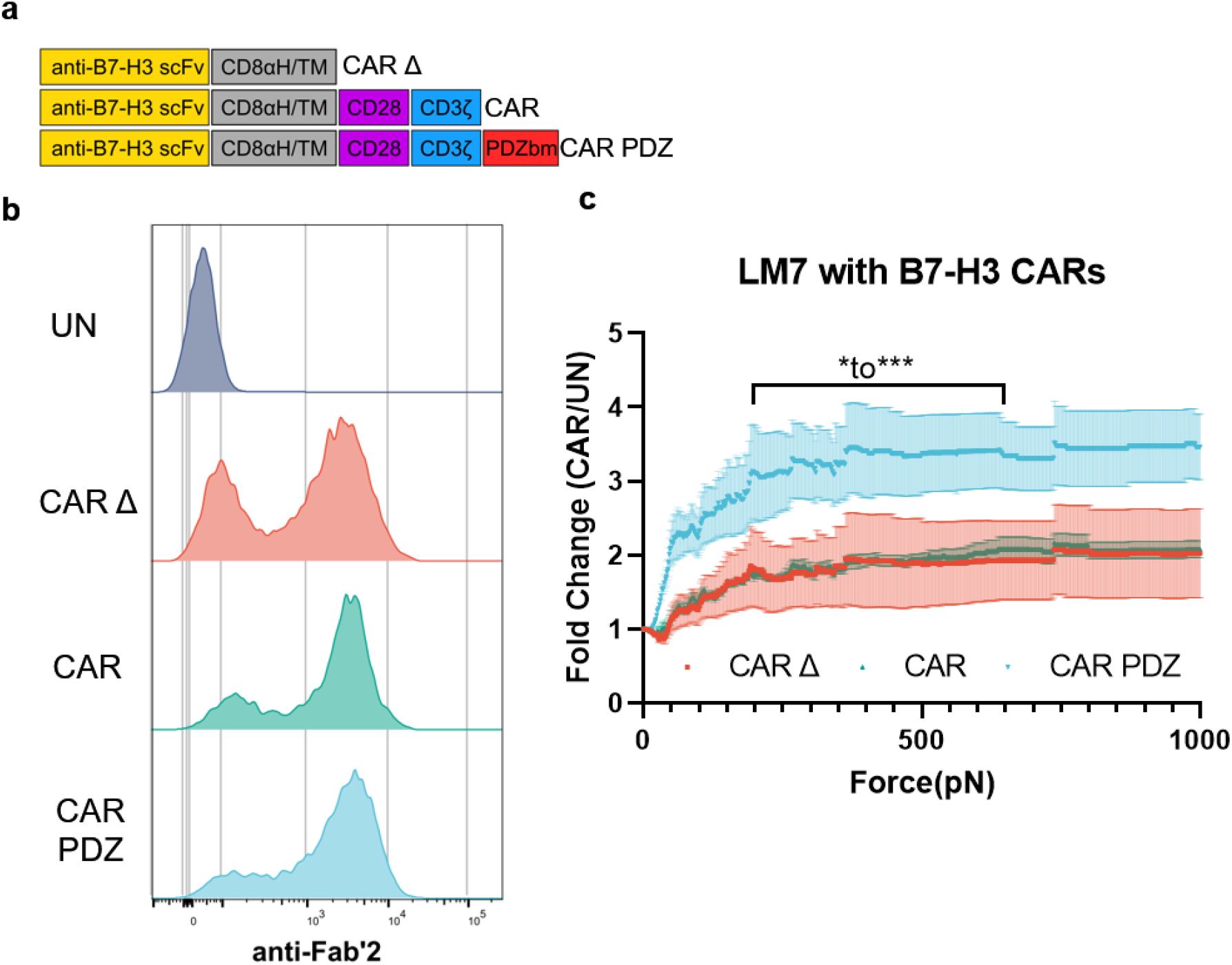
B7-H3 CAR design and Avidity Measurements. (a) Chimeric antigen receptor design schemes. Antigen recognition domain (anti-B7-H3 scFv): goldenrod, hinge and transmembrane domains (CD8αH/TM): grey, CD28 co-stimulatory domain: purple, CD3ζ activation domain: blue, PDZbm scaffolding anchor domain: red. (b) Example flow cytometry plot detailing B7-H3 CAR expression. (c) Normalized fold change of CAR NK cell binding compared to untransduced NK cells. Bracketed line indicates the scale of statistical difference at CAR.PDZ vs. CAR Δ and CAR from 194 to 646pN for both comparisons except for 205-215pN. Two-Way ANOVA was used to determine statistical significance with Two-stage linear step-up procedure of Benjamini, Krieger and Yekutieli to correct for FDR; q<0.05 *, <0.001 ***, n=3, mean±SEM shown.

**Extended Data Figure 6:**
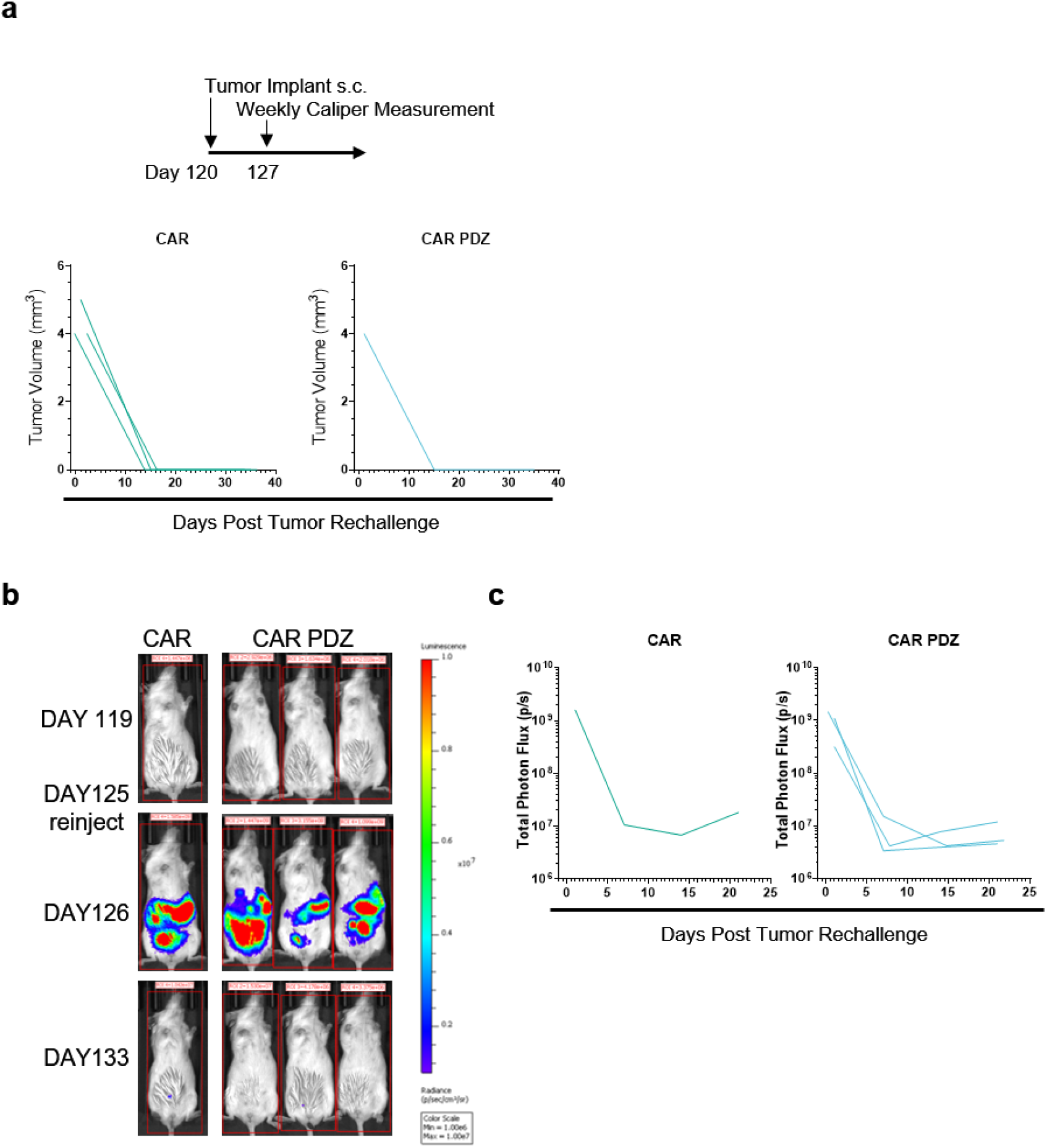
A549 and LM7 tumor rechallenge rejection. (a) A549 tumor rechallenge timeline with identical initial cancer cell numbers. Indicated tumor volumes from palpable nodules overtime. (b) Intravital imaging of LM7 rechallenge with identical initial cancer cell numbers in complete responder mice. Color scale 1e^6^ to 1e^7^ of total photon flux(p/s). (c) Tumor flux values of weekly measurements.

**Extended Data Table 1:**
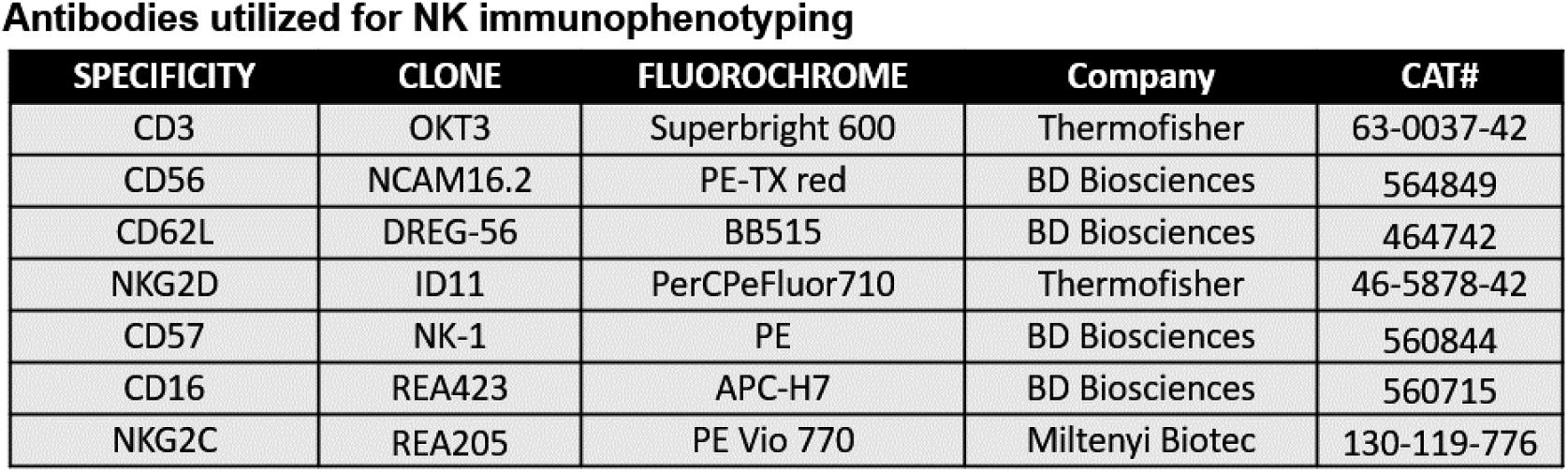
Immunophenotyping Antibody Panel.

## Notes

### Competing Interest Statement

Patents and patent applications in the immunotherapy field, G.K., P.C., and S.G. Research collaboration with TESSA Therapeutics, is a DSMB member of Immatics, and on the scientific advisory board of Tidal, S.G. Technical Consultant for LUMICKS, P.C.

